# Genomic and ecological characterization of unclassified tetraploid *Echinochloa* plants distributed in northern Japan

**DOI:** 10.64898/2026.09.08.750280

**Authors:** Tomomi Kubo, Hikari Ikeda, Kentaro Yasuda, Shunji Kurokawa

## Abstract

Understanding the evolutionary origins of globally noxious weeds provides insights into the genetic background of weediness. Among species of *Echinochloa*, the hexaploid *E. crus-galli* was previously thought to have arisen through hybridization and allopolyploidization between tetraploid *E. oryzicola* and an undiscovered diploid species, although they exhibit marked differences in both ecological and morphological characteristics. Recently, in Japan, unknown tetraploid *Echinochloa* plants were discovered that exhibit distinct ecological and morphological characteristics from those of *E. oryzicola*, with some traits showing greater similarity to those of *E. crus-galli*. In this study, we investigated the relationships between well-known *Echinochloa* species and these unknown tetraploid plants, hereafter referred to as “ninja”. Subgenome-based read mapping analysis and phylogenetic analysis revealed that ninja possesses a distinct genomic composition, while showing close relationships to *E. oryzicola* in both its chloroplast and nuclear genomes. Furthermore, the identification of a very small number of ninja × *E. oryzicola* hybrids among field-collected individuals suggests that these two species rarely hybridize in the wild. Although ninja and *E. oryzicola* showed similar genomic compositions, ninja has a broader habitat range, partially overlapping with that of *E. oryzicola* and extending into areas occupied by *E. crus-galli*. Our findings indicate the possibility that hexaploid *E. crus-galli* may have originated from a tetraploid *Echinochloa* species other than *E. oryzicola*.

## 1. INTRODUCTION

Knowledge of the evolutionary history of noxious weeds is crucial to uncover their weediness. The genus *Echinochloa* comprises approximately 50 species (Michael, 2001), including both weeds and crops, with a basic chromosome number (x) of nine and ploidy levels ranging from diploid (2x) to 14-ploid (14x) (Tanesaka, 1991; Yabuno, 1966). In particular, tetraploid *E. oryzicola* (Vasinger) Vasinger (late watergrass) and hexaploid *E. crus-galli* (L.) Beauv (barnyardgrass) are widely distributed across East and Southeast Asia (Osada, 1989; Yabuno, 1966, 2001). Hexaploid *E. crus-galli* exhibits remarkable intraspecific diversity in ecological and morphological traits and shows broad environmental adaptability, occurring in a wide range of habitats from wet paddy fields to drier grasslands (Yabuno, 1966, 2001; Yamasue et al., 1989). In contrast, tetraploid *E. oryzicola* is predominantly distributed in paddy fields and exhibits limited morphological diversity compared to *E. crus-galli* (Yabuno, 1966; Yamasue et al., 1989). Chromosome pairing at metaphase I (MI) in *E. crus-galli* × *E. oryzicola* hybrids indicated that the two genomes of *E. oryzicola* are homologous to two of the three genomes of *E. crus-galli* (Yabuno, 1966). Based on this cytological evidence, hexaploid *E. crus-galli* is thought to have arisen through hybridization and subsequent allopolyploidization between tetraploid *E. oryzicola and an* undiscovered diploid species. In the phylogenetic analysis of the chloroplast genomes of *Echinochloa* species, the individuals of *E. crus-galli* and *E. oryzicola* each formed a distinct monophyletic clade (Aoki & Yamaguchi, 2008; Hereward et al., 2016; Jiang et al., 2021; Nah et al., 2015; Perumal et al., 2016; Sebastin et al., 2019; Yamaguchi et al., 2005; Ye et al., 2014), suggesting that *E. oryzicola* was a paternal ancestor of *E. crus-galli* (Aoki & Yamaguchi, 2008).

However, in recent analyses, several varieties of *E. crus-galli* possessing chloroplast genomes similar to those of *E. oryzicola* have been discovered in Asian countries (Gao et al., 2022; Wu et al., 2022; Yasuda & Nakayama, 2019). Wu et al. (2022) described three major types of *Echinochloa* chloroplast genome as clade0, clade1 and clade2, while almost all accessions of *E. oryzicola* and some accessions of *E. crus-galli* var. crus-galli from China and Malaysia belong to clade1, whereas most of the accessions of several varieties of *E. crus-galli* belongs to clade2. Furthermore, several tetraploid plants from Hainan Province in China were classified into clade2, in addition to the American tetraploid species *E. walteri* (Wu et al., 2022). These Hainan tetraploid plants in clade2 were assumed to possess an AABB genome composition and were designated as *E. oryzicola* var. *hainanensis*, while the other *E. oryzicola* plants in clade1 were designated as *E. oryzicola* var. *oryzicola*. Given these recent comprehensive genomic analyses, the previously proposed evolutionary origin of *E. crus-galli* may need to be reconsidered.

In 2013, unclassified *Echinochloa* plants with 36 chromosomes were discovered in northern Japan (Yasuda et al., 2020). In Japan, the previously identified tetraploid *Echinochloa* species is only *E. oryzicola* var. *oryzicola,* which exhibits erect tillers mimicking rice (Barrett, 1983; Yabuno, 1966), whereas the newly discovered tetraploid plants have semi-erect tillers. Furthermore, although the habitat of tetraploid plants with semi-erecting tillers partially overlaps with that of *E. oryzicola* var. *oryzicola,* they tend to co-occur more frequently with *E. crus-galli*. These findings suggest that the newly discovered tetraploid *Echinochloa* plants possess a genetic background and ecological characteristics that differ from those of *E. oryzicola*. In this study, we investigated tetraploid *Echinochloa* plants of uncertain genetic classification and ecological characteristics (Yasuda et al., 2020), hereafter referred to as “ninja.” By comparing the genetic and distributional characteristics of ninja with those of previously reported *Echinochloa* species, we aimed to clarify the potential role of ninja in the establishment of *Echinochloa* weeds in farmlands. First, to determine the genetic relationship between ninja and other species, we analyzed resequencing data from representative accessions of *Echinochloa* species. We also analyzed the population structure using dpMIG-seq data from nationwide samples collected in the 1970s and field-collected samples from northern Japan in 2024. Furthermore, to clarify the distributional characteristics of ninja, vegetation and environmental surveys were conducted across ninja collection sites. By integrating these environmental information, we compared the habitat of ninja with that of other species.

## 2. MATERIALS & METHODS

### 2.1 Resequencing data of global *Echinochloa* species

Previously generated whole-genome resequencing data of Echinochloa accessions were obtained from the National Genomics Data Center (NGDC) under project accession numbers PRJCA003883 and PRJCA002334 for *E. crus-galli* var. *praticola*, *E. crus-galli* var. *crus-galli*, *E. crus-galli* var. *esculenta, E. colona* var. *colona*, *E. colona* var. *frumentacea, E. walteri*, *E. oryzicola* var. *hainanensis,* and *E. haploclada*. The data for *E. oryzicola* var. *oryzicola* and ninja were obtained from the DNA Data Bank of Japan (DDBJ) under project accession number PRJDB14855. The data for *Setaria italica* (L.) P. Beauv. accession was retrieved from the Sequence Read Archive (SRA) under accession number SRR3431014. These resequencing datasets comprised paired-end reads with read lengths of 100 or 150 bp (Table 1).

**Table 1.**
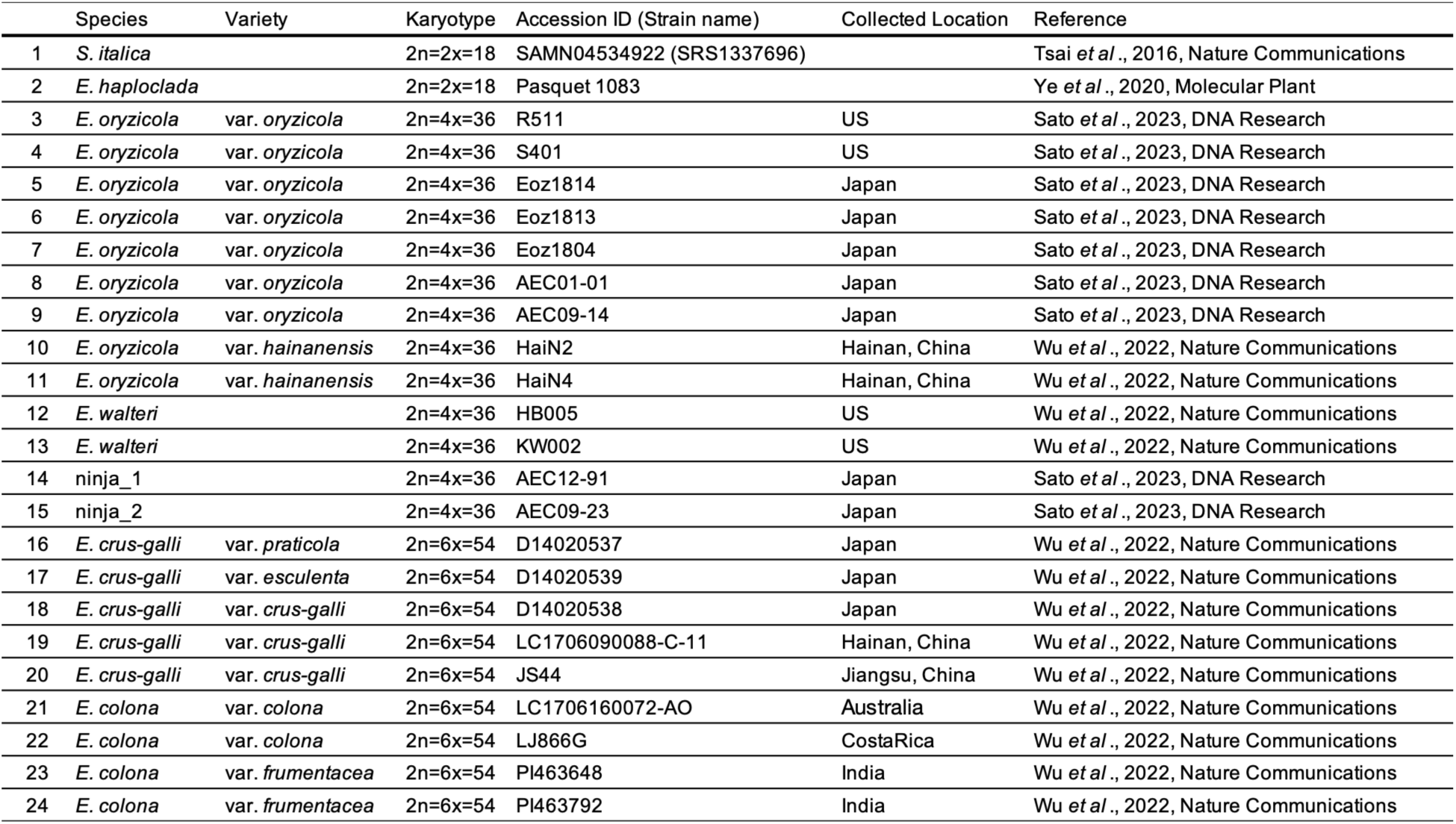
Resequencing data used for genomic analysis.

|  | Species | Variety | Karyotype | Accession ID (Strain name) | Collected Location | Reference |
| --- | --- | --- | --- | --- | --- | --- |
| 1 | *S. italica* |  | 2n=2x=18 | SAMN04534922 (SRS1337696) |  | Tsai *et al.*, 2016, Nature Communications |
| 2 | *E. haploclada* |  | 2n=2x=18 | Pasquet 1083 |  | Ye *et al.*, 2020, Molecular Plant |
| 3 | *E. oryzicola* | var. *oryzicola* | 2n=4x=36 | R511 | US | Sato *et al.*, 2023, DNA Research |
| 4 | *E. oryzicola* | var. *oryzicola* | 2n=4x=36 | S401 | US | Sato *et al.*, 2023, DNA Research |
| 5 | *E. oryzicola* | var. *oryzicola* | 2n=4x=36 | Eoz1814 | Japan | Sato *et al.*, 2023, DNA Research |
| 6 | *E. oryzicola* | var. *oryzicola* | 2n=4x=36 | Eoz1813 | Japan | Sato *et al.*, 2023, DNA Research |
| 7 | *E. oryzicola* | var. *oryzicola* | 2n=4x=36 | Eoz1804 | Japan | Sato *et al.*, 2023, DNA Research |
| 8 | *E. oryzicola* | var. *oryzicola* | 2n=4x=36 | AEC01-01 | Japan | Sato *et al.*, 2023, DNA Research |
| 9 | *E. oryzicola* | var. *oryzicola* | 2n=4x=36 | AEC09-14 | Japan | Sato *et al.*, 2023, DNA Research |
| 10 | *E. oryzicola* | var. *hainanensis* | 2n=4x=36 | HaiN2 | Hainan, China | Wu *et al.*, 2022, Nature Communications |
| 11 | *E. oryzicola* | var. *hainanensis* | 2n=4x=36 | HaiN4 | Hainan, China | Wu *et al.*, 2022, Nature Communications |
| 12 | *E. walteri* |  | 2n=4x=36 | HB005 | US | Wu *et al.*, 2022, Nature Communications |
| 13 | *E. walteri* |  | 2n=4x=36 | KW002 | US | Wu *et al.*, 2022, Nature Communications |
| 14 | ninja_1 |  | 2n=4x=36 | AEC12-91 | Japan | Sato *et al.*, 2023, DNA Research |
| 15 | ninja_2 |  | 2n=4x=36 | AEC09-23 | Japan | Sato *et al.*, 2023, DNA Research |
| 16 | *E. crus-galli* | var. *praticola* | 2n=6x=54 | D14020537 | Japan | Wu *et al.*, 2022, Nature Communications |
| 17 | *E. crus-galli* | var. *esculenta* | 2n=6x=54 | D14020539 | Japan | Wu *et al.*, 2022, Nature Communications |
| 18 | *E. crus-galli* | var. *crus-galli* | 2n=6x=54 | D14020538 | Japan | Wu *et al.*, 2022, Nature Communications |
| 19 | *E. crus-galli* | var. *crus-galli* | 2n=6x=54 | LC1706090088-C-11 | Hainan, China | Wu *et al.*, 2022, Nature Communications |
| 20 | *E. crus-galli* | var. *crus-galli* | 2n=6x=54 | JS44 | Jiangsu, China | Wu *et al.*, 2022, Nature Communications |
| 21 | *E. colona* | var. *colona* | 2n=6x=54 | LC1706160072-AO | Australia | Wu *et al.*, 2022, Nature Communications |
| 22 | *E. colona* | var. *colona* | 2n=6x=54 | LJ866G | Costa Rica | Wu *et al.*, 2022, Nature Communications |
| 23 | *E. colona* | var. *frumentacea* | 2n=6x=54 | PI463648 | India | Wu *et al.*, 2022, Nature Communications |
| 24 | *E. colona* | var. *frumentacea* | 2n=6x=54 | PI463792 | India | Wu *et al.*, 2022, Nature Communications |

### 2.2 Field collection and dpMIG-seq data generation of *Echinochloa* plants

Seeds and fresh leaves of *Echinochloa* plants were collected from 12 locations in northern Japan, where ninja had previously been identified (Yasuda et al., 2020). In September 2024, individuals were collected from paddy fields, levees, roadsides, and grasslands at densities of 0.1–1 plant m⁻¹, with 6–53 individuals from each location (Fig. 4B). In addition to the 333 field-collected individuals, standard accessions of *E. oryzicola* var. *oryzicola* (two accessions), *E. crus-galli* var. *formosensis* (three accessions), ninja_1, and ninja_2, which had previously been identified based on chromosome number and morphological characteristics (Yasuda et al., 2020; Yasuda & Nakayama, 2016), were cultivated to collect fresh leaves for further analysis. In addition, 38 accessions collected throughout Japan in the 1970s and maintained at the NARO Genebank (Genetic Resources Research Center, National Agriculture and Food Research Organization [NARO], Japan) as *E. oryzicola* var. *oryzicola*, with accession numbers JP3135–JP3567 and JP3569 were also included. Genomic DNA was extracted using the CTAB method (Murray & Thompson, 1980). dpMIG-seq libraries were prepared (Nishimura et al., 2024), and 150-bp paired-end reads were generated using the DNBSEQ-G400 platform (BGI, Shenzhen, China).

### 2.3 Reads cleaning and mapping

Paired-end whole-genome resequencing reads were subjected to quality control. Adapter sequences were removed using Cutadapt (v2.8) (Martin, 2011) or fastp (v1.0.1) (Chen, 2025) according to the preprocessing requirements of each dataset. Quality trimming was subsequently performed using Trimmomatic (v0.38) (Bolger et al., 2014) with the following settings: LEADING:25, TRAILING:25, and SLIDINGWINDOW:4:15. Reads shorter than 90 and 100 bp were discarded from the datasets initially generated with 100-bp and 150-bp reads, respectively. Read quality was assessed using FastQC (v0.11.9) (Andrews, 2010).

The 150-bp paired-end dpMIG-seq reads were processed according to a previously described workflow (Nishimura et al., 2024) with minor modifications. Adapter sequences and the first 17 bp of each read were removed using Trimmomatic (v0.38), followed by quality trimming with LEADING:25, TRAILING:25, and SLIDINGWINDOW:4:15. Reads shorter than 120 bp were discarded. Paired-end read correction was performed using fastp (v0.23.4), and quality control reports generated by fastp were aggregated using MultiQC (v1.9) (Ewels et al., 2016). Samples with unsuccessful DNA extraction and substantially low read counts were excluded. Finally, reads for 328 samples collected in 2024, 37 samples collected in the 1970s, and two or three standard accessions for each *Echinochloa* species were subjected to subsequent analyses, resulting in 372 samples.

Quality-filtered reads from both the resequencing and dpMIG-seq datasets were aligned to the corresponding reference genome using SNAP aligner (v2.0.1) (Zaharia et al., 2011). For the resequencing data, *E. oryzicola* var. *oryzicola* nuclear genome (Sato et al., 2023), *E. crus-galli* var.

*crus-galli* chloroplast genome (Ye et al., 2014) and pseudo-hexadecaploid genome “ATBT + AHBHCH + DHEHFH,” which integrates previously reported *Echinochloa* genome sequences (Guo et al., 2017; Sato et al., 2023; Wu et al., 2022; Ye et al., 2020) were used as references. For the dpMIG-seq data, reads were mapped independently to the *E. oryzicola* var. *oryzicola* nuclear genome and *E. crus-galli* var. *crus-galli* nuclear genome. The alignment parameters were consistent among the samples within each dataset and read group information was assigned during alignment. Secondary alignments were excluded, and only primary alignments were retained for further analyses. The resulting BAM files were indexed using SAMtools (v1.9) (Li et al., 2009). Mapped reads were subsequently filtered with bamutils from ngsutils (v0.5.9) (Breese & Liu, 2013), retaining properly paired alignments with no more than three mismatches relative to the reference sequences. For standard accessions in the dpMIG-seq datasets, technical replicate libraries generated from the same DNA templates were combined after BAM filtering to increase the sequencing depth for key reference accessions using SAMtools (v1.9), resulting in approximately twice the read depth of the field-collected samples.

### 2.4 Mapping-based analysis of genome composition and ploidy levels

Subgenome-specific read mapping patterns were analyzed to infer the genome composition and ploidy levels of the *Echinochloa* samples using BAM files generated from both whole-genome resequencing and dpMIG-seq datasets. BAM indexing and read-count statistics were performed using SAMtools (v1.9), whereas SAMtools (v1.19.2) was used for breadth-of-coverage analysis. Specifically, samtools stats and samtools idxstats were used to obtain the total number of sequenced reads, number of mapped reads, and number of reads mapped to individual subgenomic groups. The breadth of coverage of each subgenomic region was calculated using samtools coverage by summing the covered bases and reference sequence lengths across the chromosomes assigned to each subgenome. Statistical analyses and data visualization were performed using R (v4.4.2) (R Core Team, 2024).

For the whole-genome resequencing dataset, BAM files generated by mapping reads to a pseudo-hexadecaploid reference genome were analyzed to characterize the distribution of mapped reads among the subgenomes. The number of mapped reads assigned to each subgenome was extracted and expressed as a percentage of the total number of mapped reads. The breadth of coverage was calculated independently for each subgenome. Overall mapping rates and subgenome-specific read mapping proportions were compared among species to characterize the genomic composition of the target accession, including ninja.

For the dpMIG-seq dataset, BAM files generated by mapping reads to the *E. crus-galli* var. *crus-galli* nuclear genome were analyzed to assess the presence and relative representation of the three subgenomes (A, B, and C). The number of mapped reads assigned to each subgenome was then calculated. The C-subgenome mapping ratio was expressed as the percentage of reads assigned to the C subgenome relative to the total number of reads assigned to all three subgenomes.

The C-subgenome mapping ratios were compared between the standard accessions with known ploidy levels. Based on the observed separation between the standard tetraploid (4x) and hexaploid (6x) accessions (Fig. 3), samples with a C-subgenome mapping ratio of <15% were classified as putative 4x, whereas those with a ratio of ≥15% were classified as putative 6x. The distribution of mapped reads across the A, B, and C subgenomes was examined to evaluate the consistency of the ploidy assignments.

### 2.5 Variant calling and SNP filtering

Filtered BAM files from both the resequencing and dpMIG-seq datasets were processed with elprep (v5.1.3) (Herzeel et al., 2015) for coordinate sorting and variant calling using the HaplotypeCaller mode. Variant calling was performed independently for each reference genome dataset. The resulting gVCF files were indexed using Tabix (v1.11) (Li, 2011). Joint genotyping across samples was performed using GATK (v4.2.6.1) (Mckenna et al., 2010). Individual gVCF files were imported into GenomicsDB using GenomicsDBImport, followed by genotype inference using GenotypeGVCFs.

Using GATK (v4.2.6.1), only biallelic SNPs were retained, and genotype calls with a read depth (DP) of zero were converted to missing data (./.). Variant-level quality filtering was then applied using the same criteria for both resequencing and dpMIG-seq datasets. Variants were excluded if they met any of the following conditions: Quality by Depth (QD) < 2.0, variant quality (QUAL) < 30.0, Strand Odds Ratio (SOR) > 4.0, Fisher Strand bias (FS) > 60.0, or Root Mean Square Mapping Quality (MQ) < 40.0. Furthermore, variants with more than 20% missing genotypes (max-missing < 0.8) or a mean sequencing depth below five (min-meanDP < 5) were excluded using VCFtools (v0.1.16) (Danecek et al., 2011). Variants with a minor allele frequency (MAF) below 0.05 were removed from resequencing datasets. For the dpMIG-seq dataset, an MAF threshold of 0.01 was used to retain low-frequency variants that could be informative for identifying genetically differentiated minor lineages within the Japanese *Echinochloa* population.

### 2.6 Phylogenetic analysis

Filtered SNPs were converted from VCF to FASTA format using Python (v3.12.3) (*Python*, 2024). Missing genotypes were represented as N, whereas heterozygous genotypes were encoded using the corresponding IUPAC ambiguity codes. To accommodate the ascertainment bias correction implemented in RAxML (v8.2.13) (Stamatakis, 2014), only polymorphic sites for which at least two nucleotide states observed in homozygous genotypes were retained. For the resequencing datasets, 586 and 10,053,414 SNPs were retained for the chloroplast and nuclear genome datasets, respectively. For the dpMIG-seq dataset, 24,012 SNPs were retained in the nuclear genome dataset.

Maximum-likelihood phylogenetic analyses were performed using RAxML (v8.2.13) under the ASC_GTRGAMMA substitution model with Lewis ascertainment bias correction. For the resequencing chloroplast genome dataset, *S. italica* was designated as the outgroup, and branch support was assessed using 1,000 rapid bootstrap replicates while simultaneously searching for the best-scoring maximum likelihood tree. For the resequencing nuclear genome dataset, *E. walteri* was used as the outgroup, and the same analytical procedure was performed using 100 rapid bootstrap replicates. For the dpMIG-seq nuclear genome dataset, the standard accession of *E. crus-galli* var. *formosensis* was designated as the outgroup, and node support was assessed using 1,000 rapid bootstrap replicates. The resulting phylogenetic trees were visualized using the Interactive Tree of Life (Letunic & Bork, 2024).

### 2.7 Population structure analysis

Population structure analysis was performed using nuclear genome-derived SNP datasets. To reduce the influence of linkage disequilibrium (LD) among closely located SNPs, LD pruning was conducted on the quality-filtered SNP dataset using PLINK (v1.90b6.21) (Chang et al., 2015). SNPs were pruned using a sliding window approach with a window size of 50 SNPs, a step size of 10 SNPs, and a r² threshold of 0.5.

Principal component analysis (PCA) was subsequently performed using the LD-pruned SNP dataset with the --pca option implemented in PLINK. The proportion of variance explained by each principal component was calculated based on the eigenvalues generated using the PLINK software. PCA plots were generated using R.

Individual ancestry proportions were estimated using ADMIXTURE (v1.3.0) (Alexander et al., 2009). ADMIXTURE analyses were performed for K = 1–15 ancestral clusters, with cross-validation enabled to evaluate the model’s fit across K values. The resulting cross-validation errors were compared among the different K values. ADMIXTURE results were processed and visualized in R, and the ancestry proportions for K = 1–7 are presented in the final plots. Individuals were arranged according to the PCA-defined genetic group.

### 2.8 Hybrid Index analysis

To evaluate the possibility of hybridization between different genetic groups, we calculated the hybrid index (HI) and interclass heterozygosity to create a triangle plot (Fitzpatrick, 2012; Wiens et al., 2025). SNPs with an allele frequency difference >0.9 between each pair of putative parental groups were identified as ancestry-informative markers (AIMs), and HI and heterozygosity were calculated for each individual based on these markers. The resulting values were visualized in triangle plots using R and compared with the expected distributions of the parental groups.

### 2.9 Environmental survey of ninja habitat

To characterize the habitat conditions of ninja, vegetation and environmental surveys were conducted in September 2025 at a subset of the collection sites in 2024 (Sites 1–6; Fig. 4, Fig. S5). At each site, two to six 50 × 50 cm quadrats were established to include *Echinochloa* plants. Within each quadrat, total vegetation cover, plant species, species-specific cover, and plant height were recorded, and soil samples were collected. Plant species were identified based on their morphological characteristics. To quantify the wetness and light conditions at the survey sites, a digital elevation model (DEM) was constructed in QGIS (v3.40.11) (Dawson et al., 2025) using the Fundamental Geospatial Data (Digital Elevation Model; 5- and 10-m meshes) provided by the Geospatial Information Authority of Japan, and the missing values were interpolated. Based on the DEM, flow accumulation and slope were calculated to derive the Topographic Wetness Index (TWI), while the total incoming solar radiation for June 1, 2025, was estimated using the Potential Incoming Solar Radiation tool in SAGA GIS (v7.8.2) (Conrad et al., 2015). The raster values of the TWI and solar radiation were then extracted for each survey site. Collected soil samples were air-dried, passed through a 2,000-µm sieve, and analyzed for their chemical and physical properties (Tokachi Federation of Agricultural Cooperatives, Hokkaido, Japan).

To characterize the vegetational composition and environmental conditions among quadrats, PCA was performed by integrating vegetation and environmental data. Species-specific dominance values were calculated as the product of species-specific cover and plant height, converted to relative dominance based on the total dominance within each quadrat, and subjected to arcsine square-root transformation. The transformed values were used as vegetation variables. The environmental variables included soil properties, TWI, and potential incoming solar radiation. The following soil variables were used as soil properties: soil pH (H₂O), available phosphorus (P), exchangeable potassium (K), exchangeable magnesium (Mg), exchangeable calcium (Ca), Mg/K ratio, Ca/Mg ratio, Ca saturation (%), base saturation (%), total nitrogen (%), nitrate nitrogen (NO₃⁻–N), ammonium nitrogen (NH₄⁺–N), phosphate absorption coefficient, cation exchange capacity (CEC), and bulk density. The percentage values for Ca saturation, base saturation, and total nitrogen were arcsine square-root transformed. As soil samples could not be obtained from quadrats 4–6 at Site 2, the mean values of quadrats 1–3 at the same site were used. All vegetative and environmental variables were standardized, and those with no variance were removed. PCA was performed using the prcomp function in R.

## 3 RESULTS

### 3.1 Mapping analysis to a pseudo-hexadecaploid reference genome

First, the subgenome composition of ninja was compared with that of other *Echinochloa* species. Resequencing reads were mapped to a pseudo-hexadecaploid genome, “ATBT + AHBHCH + DHEHFH,” which integrates previously reported reference genome sequences of *Echinochloa* species. Subgenome-specific genome coverages and mapping rates were compared among species to determine the relative mapping preferences for each subgenome. In this analysis, multimapped reads were excluded, and only uniquely mapped reads were counted, accounting for approximately 50–75% of the total reads (Fig. S1A). Reads from *E. oryzicola* var. *oryzicola*, *E. crus-galli*, and *E. colona*, which constitute the reference genome sequences, were mapped to the corresponding reference subgenomes (Fig. 1, Fig. S1B). This result indicates that the subgenome composition of each sample is correctly reflected in the breadth of genome coverage and read-mapping proportions when mapped to the pseudo-hexadecaploid genome sequence. Reads from the ninja were mapped to the “ATBT” and “AHBH” subgenomes but not to the “CH” or “DHEHFH” subgenomes, similar to those from the tetraploid *E. walteri* inhabiting America and *E. oryzicola* var. *hainanensis* in China. Therefore, in comparison with previously reported subgenome compositions, it is inferred that ninja possesses subgenomes similar to the ATBT of *E. oryzicola* var. *oryzicola* and AHBH of *E. crus-galli*.

**Figure 1.**
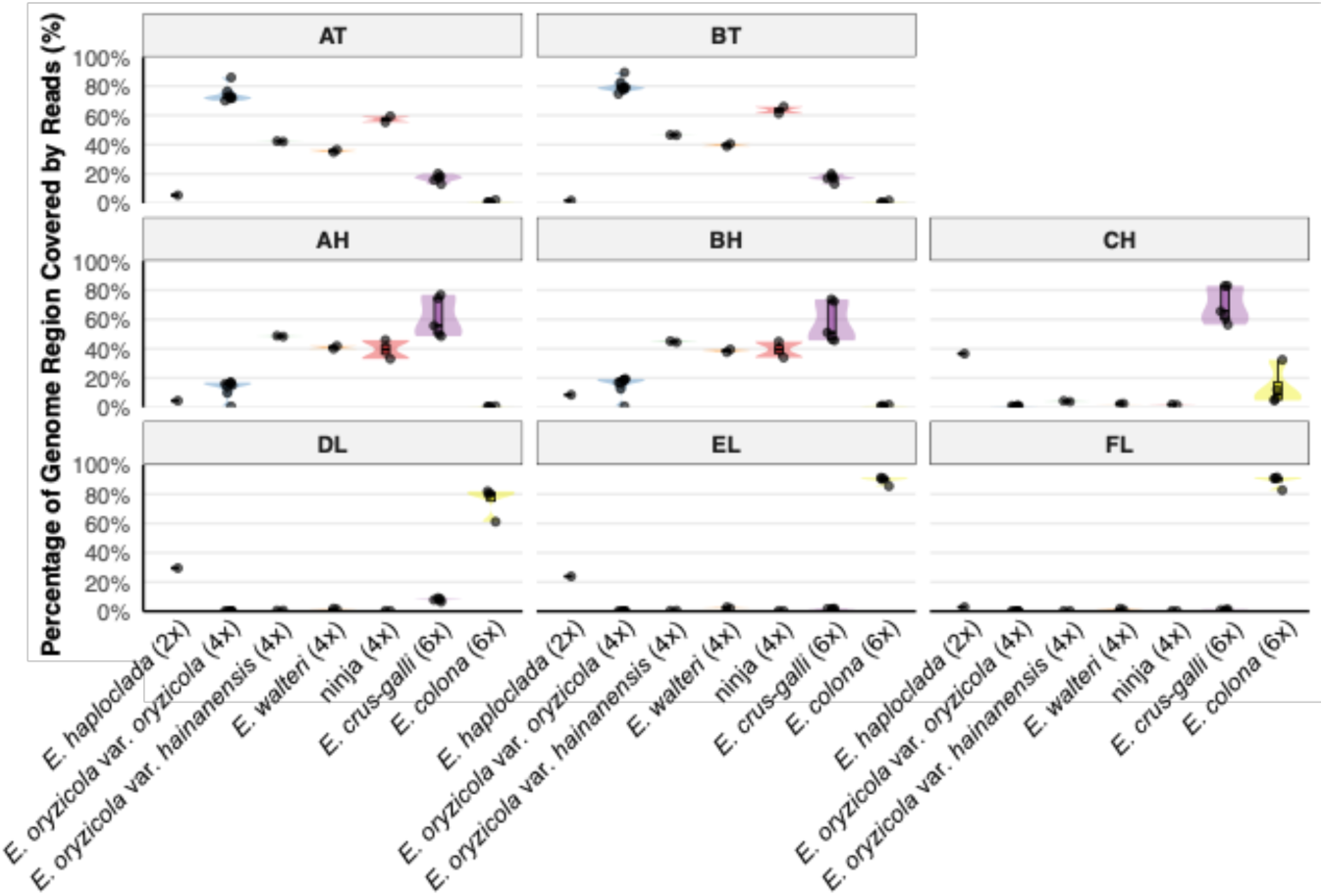
Breadth of genome coverage of the pseudo-hexadecaploid genome. The resequencing reads of *Echinochloa species,* including ninja, were mapped to a pseudo-hexadecaploid genome comprising ATBT, AHBHCH, and DHEHFH to compare their genomic compositions. The breadth of genome coverage was compared among species.

### 3.2 Phylogenetic analysis of chloroplast genomes

In the phylogenetic tree of the chloroplast genomes, the closely related Poaceae species *S. italica was* used as the outgroup (Gao et al., 2022; Guo et al., 2017), and *E. colona* var. *colona* and *E. colona* var. *frumentacea* branched off immediately, forming a monophyletic group at the root (ECpG_clade0). The other *Echinochloa* species were divided into two clades, “ECpG_clade1” constituting tetraploid *E. oryzicola* var. *oryzicola*, ninja, and hexaploid *E. crus-galli* var. *crus-galli* or “ECpG_clade2” constituting deploid *E. haploclada*, tetraploid *E. walteri,* and *E. oryzicola* var. *hainanensis,* and hexaploid *E. crus-galli* var. *crusgalli, E. crus-galli* var. *esculenta* and *E. crus-galli* var. *praticola* (Fig. 2A). ECp_clade1 corresponds to clade1 and ECpG_clade2 to clade2 in a previous study (Wu et al., 2022). In ECpG_clade1, ninja did not form a monophyletic group. Among the ninja accessions, ninja_1 was closely related to *E. crus-galli* in ECpG_clade1. In contrast, ninja_2 was closely related to *E. oryzicola* var. *oryzicola*. These findings suggest that ninja diverged multiple times in ECpG_clade1.

**Figure 2.**
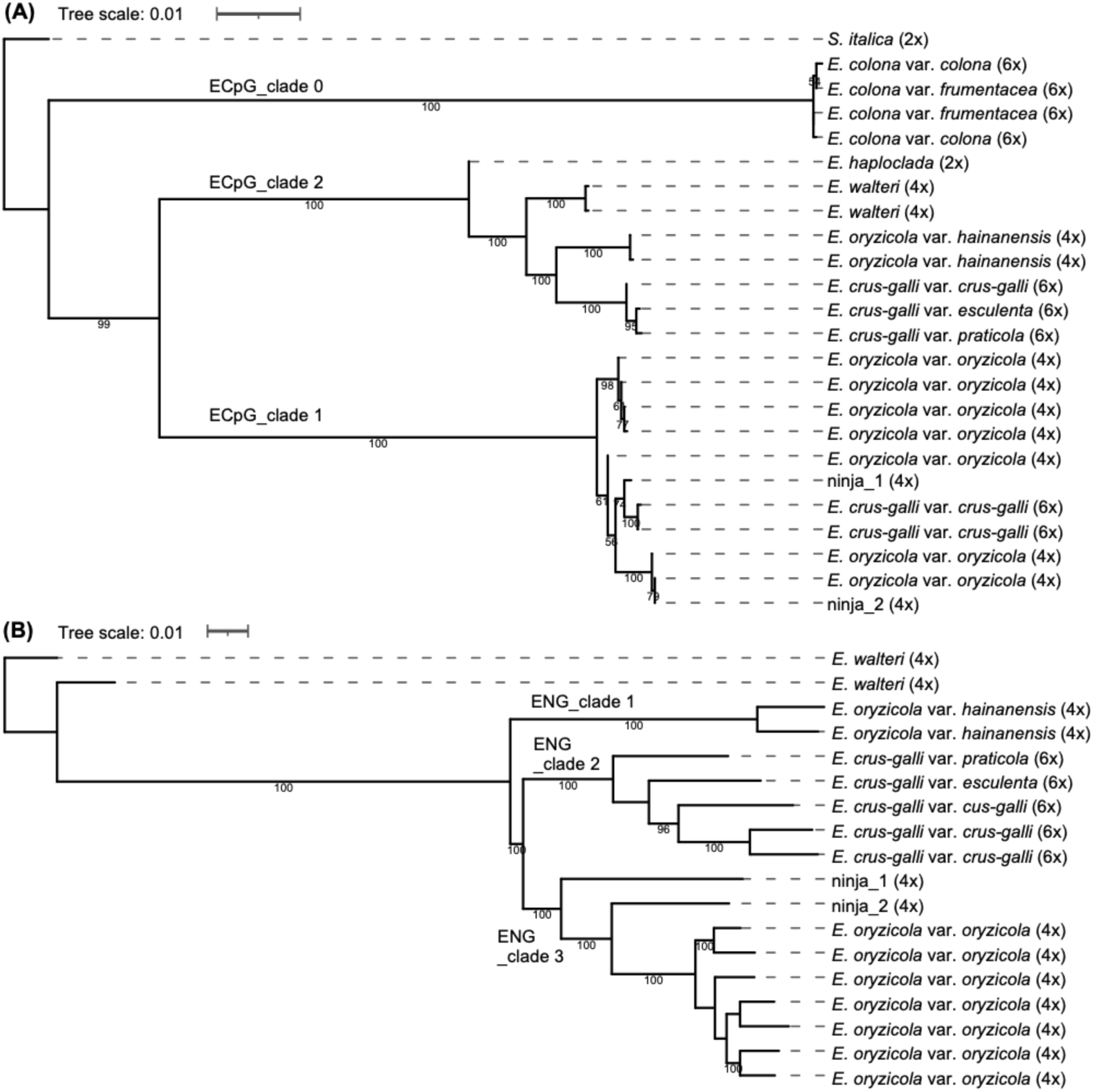
Phylogenetic tree of the genus *Echinochloa*. **A: Chloroplast phylogeny** **B: Nuclear genome (AABB) phylogeny** The resequencing reads were mapped to the chloroplast genome of E. crus-galli and the nuclear genome of E. oryzicola. Nuclear-derived SNPs were used to infer the phylogenetic relationships between ninja and other Echinochloa species. Bootstrap values ≥50 are shown on the tree.

### 3.3 Phylogenetic analysis and PCA of nuclear genomes

For *Echinochloa* species sharing an AABB-like subgenome composition, the nuclear phylogeny was analyzed using *E. oryzicola* var. *oryzicola* as the reference sequence and *E. walteri* as an outgroup. In the nuclear phylogeny, all accessions of tetraploid *E. oryzicola* var. *hainanensis* branched off immediately, forming a monophyletic group (ENG_clade1), and the remaining *Echinochloa* species were separated into hexaploid *E. crus-galli* (ENG_clade2) and tetraploid *E. oryzicola* var. *oryzicola,* and ninja (ENG_clade3) (Fig. 2B). Within ENG_clade3, ninja did not form a monophyletic group, branching in the order of ninja_1, ninja_2, and *E. oryzicola* var. *oryzicola*. In the PCA plot based on nuclear genome sequences, ninja exhibited a genetic composition similar to that of *E. oryzicola* var. *oryzicola* among tetraploid *Echinochloa* species (Fig. S2). However, the plots for the two ninja accessions were more dispersed than those for *E. oryzicola* var. *oryzicola*, which contains both Japanese and American accessions, suggesting that the two ninja accessions have distinct genetic compositions.

### 3.4 Ploidy analysis of the collected samples

To explore the distribution and population structure of *Echinochloa* species, including ninja, 333 individuals were collected from 12 locations in northern Japan where ninja had previously been identified. In addition, standard accessions of ninja, *E. oryzicola* var. *oryzicola*, and *E. crus-galli*, as well as 38 accessions collected across Japan in the 1970s and preserved as *E. oryzicola* var. *oryzicola* were subjected to genomic investigation. First, the dpMIG-seq reads obtained from each sample were mapped to the reference sequence of the *E. crus-galli* nuclear genome, and the ploidy of the collected samples was determined based on the presence or absence of reads derived from the C subgenome of *E. crus-galli*. Read data from both the tetraploid (4x) and hexaploid (6x) standard accessions were mapped to the A and B subgenomes, whereas only the read data from the 6x standard accessions were mapped to the C subgenome (Fig. 3A). When the proportion of mapped reads to the C subgenome was calculated for all mapped reads, the standard accessions from each ploidy showed a clear separation, with 4x standards at approximately 5% and 6x standards at approximately 30% (Fig. 3B). From this result, it was considered possible to determine the ploidy of the unanalyzed samples using the mapping ratio to the C subgenome, with a threshold of less than 15% (4x) or greater than 15% (6x). The C subgenome mapping ratio for field-collected samples in 2024 and in the 1970s with unknown ploidy were separated into two groups, approximately 5% and 30% (Fig. 3B). Based on the ploidy criterion, they were subsequently designated as “putative 4x samples” and “putative 6x samples.”

**Figure 3.**
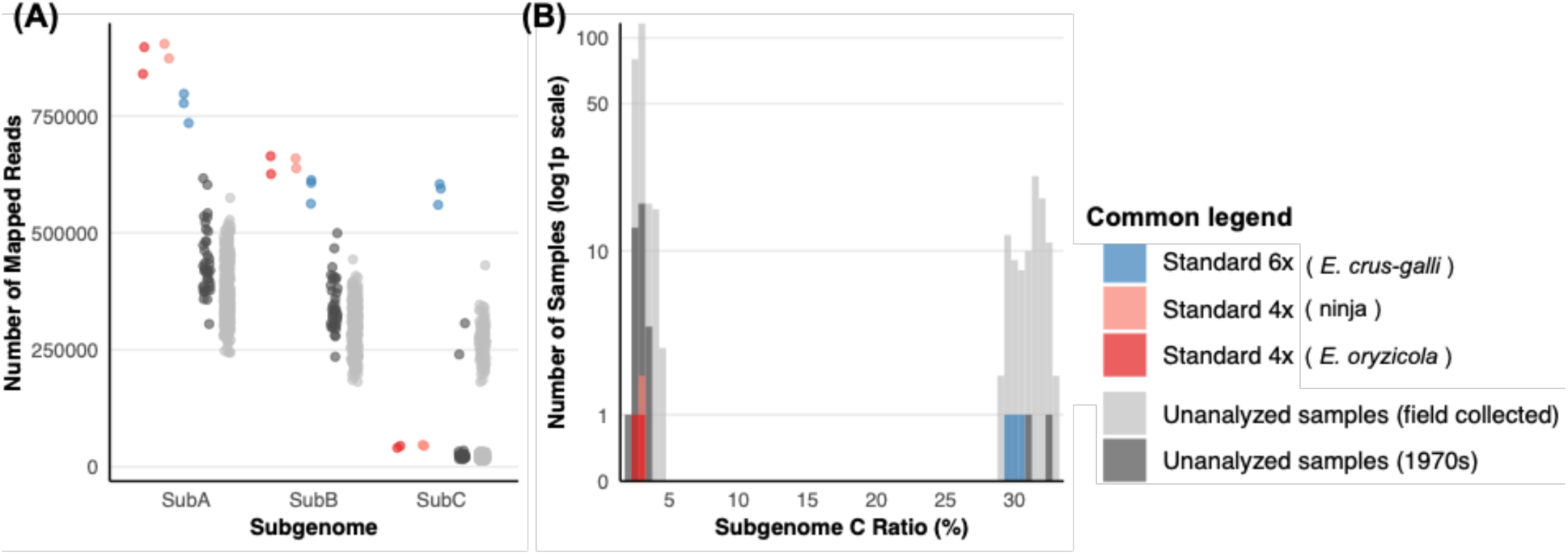
Ploidy analysis of the collected samples. **A: Mapped reads distribution to each subgenome** **B: Subgenome C mapping ratio** The dpMIG-seq data from tetraploid (4x) and hexaploid (6x) standard accessions, as well as field-collected samples from the 1970s and 2024, were mapped to the nuclear genome sequence of *E. crus-galli* (AHBHCH). For each standard accession, replicate dpMIG-seq libraries were prepared, resulting in approximately twice the read depth of the field-collected samples. The number of reads mapped to each subgenome (A) and the proportion mapped to subgenome C (B) are shown.

### 3.5 Population structure analysis of the collected samples

Next, to compare the genetic composition of each sample, SNPs were extracted using the nuclear sequence of *E. oryzicola* var. *oryzicola* as a reference, and PCA was performed. On the PC1 axis, the standard accessions were divided into 6x and 4x species, whereas on the PC2 axis, the 4x species *E. oryzicola* var. *oryzicola*, ninja_1, and ninja_2 were separated (Fig. 4A), indicating that each standard accession of ninja possesses a distinct genetic composition compared with that of *E. oryzicola* var. *oryzicola*. The 2024 field-collected samples and preserved accessions collected in the 1970s formed clusters with each standard accession. Together with the results of ploidy analysis, these samples were classified into five groups: Group1, composed of putative 4x samples and *E. oryzicola* var. *oryzicola* standards; Group2, of putative 4x samples and ninja_1 standard; Group3, of putative 4x samples and ninja_2 standard; Group4, consisting only of putative 4x samples from the 2024 field collection; and Group5, of putative 6x samples and *E. crus-galli* standard. These results demonstrate that there are at least five genetic groups of *Echinochloa* plants in northern Japan. Because Group2 and Group4 formed clusters in intermediate positions among multiple groups, it was suggested that they possessed genetic compositions intermediate to those of the other groups. Additionally, as some samples collected in the 1970s were classified in Group3, which shows a genetic composition similar to the ninja_2 standard, it was revealed that the ninja_2 like *Echinohcloa* plants were already present in Japan in the 1970s. The proportion of each group at the collecting sites confirmed that *Echinochloa* plants belonging to Group2 and Group3, which have a genetic composition similar to that of ninja, sympatrically grow with *Echinochloa* plants belonging to Group1 (including *E. oryzicola* var. *oryzicola*) and Group5 (including *E. crus-galli*) (Fig. 4B). Regarding the geographical characteristics of the distribution sites, Group1, Group3, and Group5 were widely distributed, while Group2 tended to be predominantly found on the western side of the mountain running through northern Japan. Group4 was only identified at one of the 12 sites.

**Figure 4.**
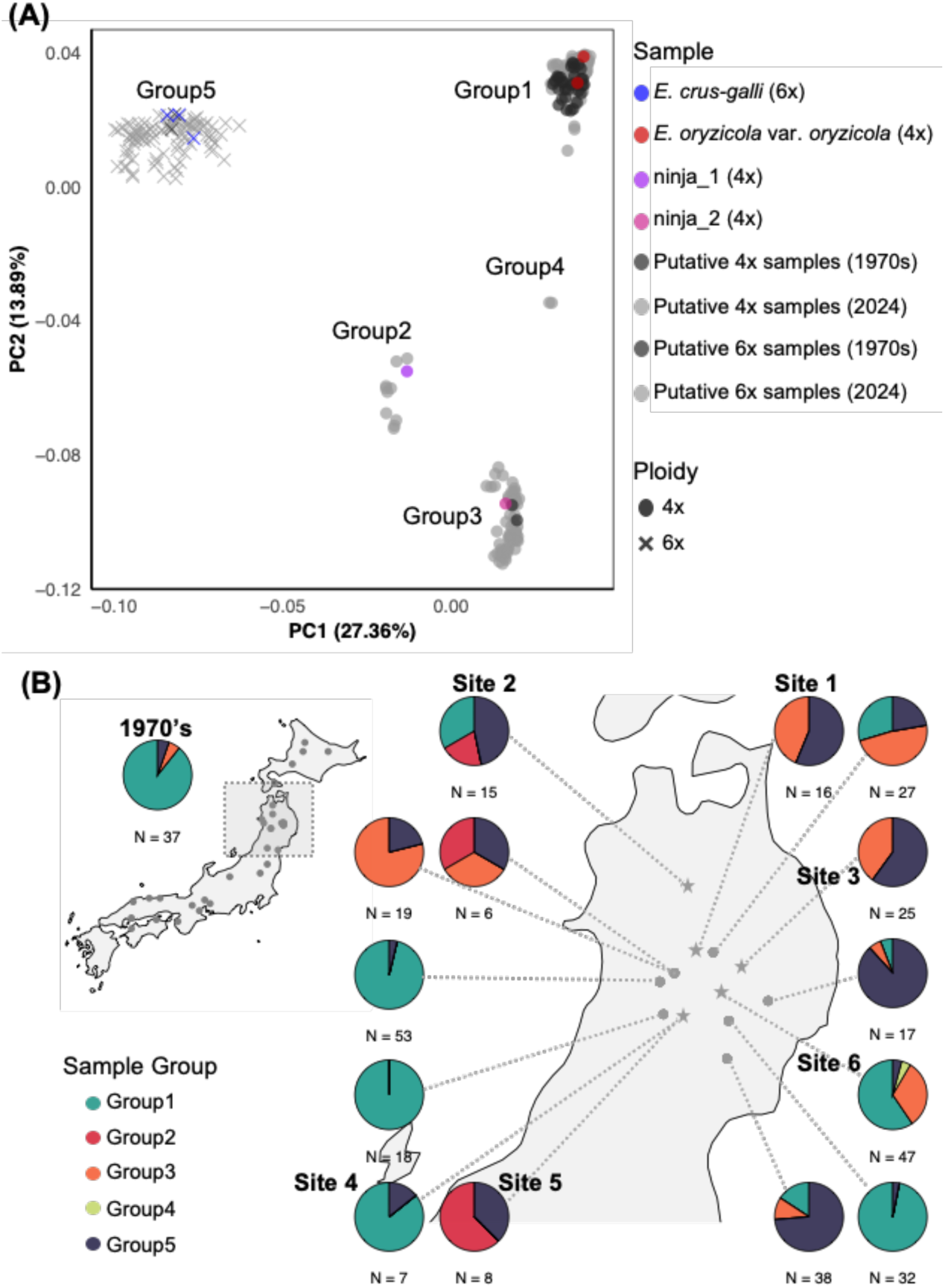
Population structure analysis of the collected samples. **A: Genomic grouping based on the principle component analysis (PCA) using SNPs derived from nuclear genome** **B: Geographical distribution and composition of the *Echinochloa* genomic groups.** The dpMIG-seq reads obtained from the collected samples were mapped to the nuclear genome of E. oryzicola (ATBT), and SNPs were identified for population structure analysis. Based on ploidy analysis and PC1 and PC2 scores from the PCA, the Echinochloa samples were classified into five groups (Groups 1–5) (A). The left map shows the sampling locations of 37 *Echinochloa* samples collected across Japan in the 1970s, and the accompanying pie chart shows the proportions of the five genomic groups among these samples. The right map shows the sampling locations in northern Japan in 2024, with pie charts showing the genomic group composition of 6–53 individuals collected from each site. Sites used for subsequent habitat condition analysis (Sites 1–6) are indicated by star symbols.

### 3.6 Phylogenetic analysis of collected samples

Next, the nuclear phylogeny of the collected samples was analyzed using a standard accession of *E. crus-galli* as an outgroup. In the phylogenetic tree, branching occurred in the order of Group5 (including the outgroup) → Group2 → Group3 → Group1 (Fig. 4C). While Group1, Group2, and Group3 each formed monophyletic group, Group4 and Group5 did not. Within the Group-forming clades, each individual generally clustered according to its collection site. Clustering by collection site was less evident in Group 5, which is considered to reflect the presence of multiple varieties of *E. crus-galli* within the group. The nuclear phylogeny suggests that Group2 and Group3, which include ninja, are not recently derived variants of Group1, which includes *E. oryzicola* var. *oryzicola* but have been maintained through independent evolutionary processes. The divergence of Group2 between Group5 and Group3, as well as the divergence of Group4 between Group3 and Group1, was consistent with the PCA results.

### 3.7 ADMIXTURE analysis of the collected samples

ADMIXTURE analysis was performed on the collected samples to visualize genetic proximity and the degree of admixture. Based on the changes in the CV error values for K=1–15, it was suggested that the model fit improved from K=3 onward (Fig. S3). Therefore, the results for K=1–7 are illustrated (Fig. 5). From K=3, where the CV error values decreased markedly, Group2 showed genetic admixture between Group3 and Group5, and Group4 showed admixture between Group1 and Group3, which was consistent with the results of the PCA and phylogenetic analysis. In addition, some Group1 individuals co-occurring with Group4 contained a small proportion of the ancestral component common to Group3. In Group5, a second component emerged from K=5, and these samples thereafter showed a distinct genetic composition from the others, which was considered to reflect genetic differences among *E. crus-galli* varieties within Group5.

**Figure 5.**
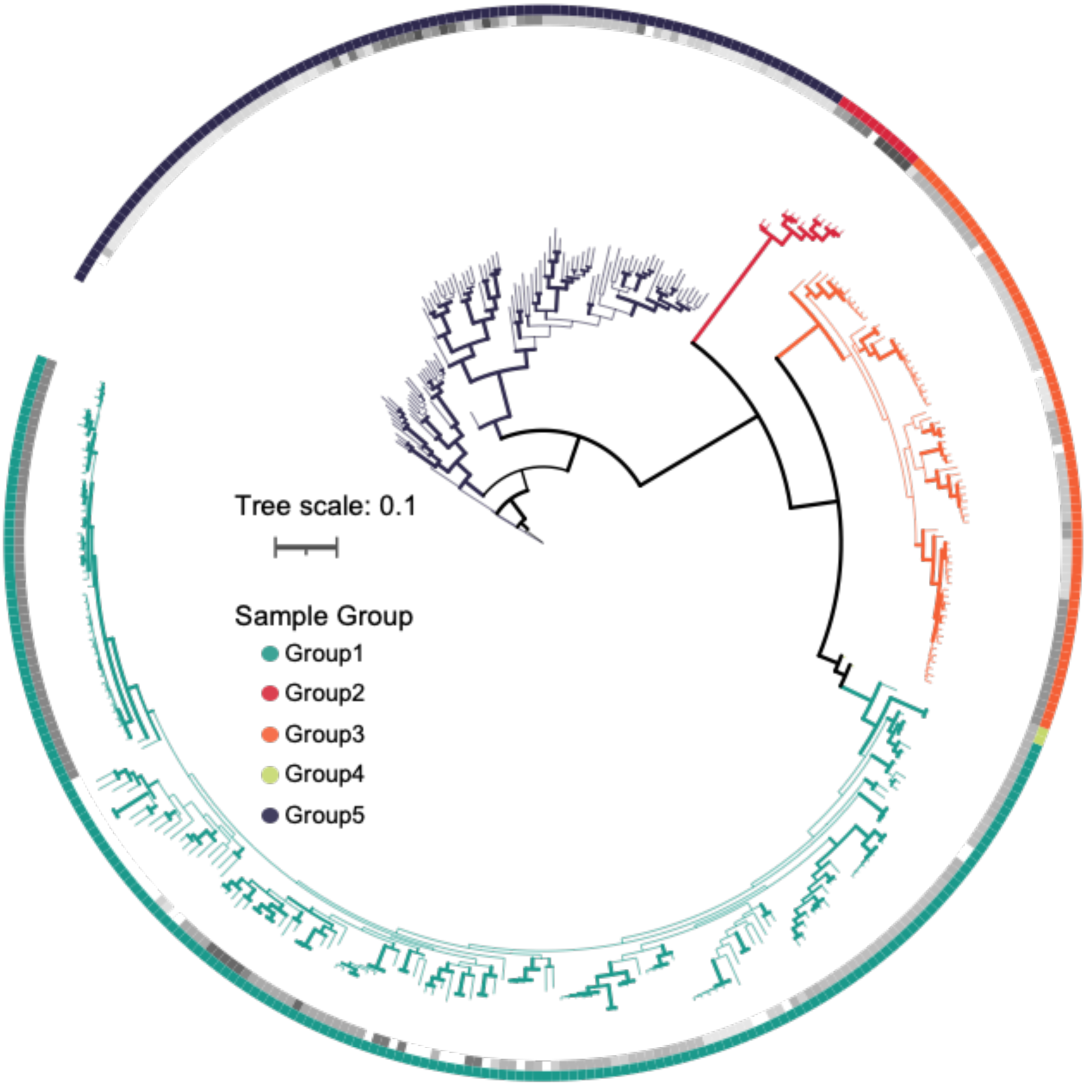
Nuclear phylogeny of the collected samples. The phylogenetic tree was constructed based on SNPs identified from the nuclear genome (ATBT) using the dpMIG-seq data from the collected samples. Bootstrap values ≥50 are indicated by branch width. The outer ring indicates the genetic group of each sample, with different colors representing the distinct groups. The inner ring indicates the sampling locations using grayscale gradients. Samples collected in the 1970s and standard accessions are shown in white in the inner ring.

### 3.8 Hybrid Index analysis in collected samples

Next, we examined the possibility that Group2 and Group4 were hybrid offspring resulting from crossing among multiple groups. We designated genomic sites with allele frequency differences of ≥ 0.9 between the putative parental groups as AIMs and investigated the HI and heterozygosity at AIMs in each sample. Group4 exhibited an intermediate HI and high heterozygosity at AIMs between Group1 and Group3 (Fig. 6), suggesting that Group4 originated from the hybridization of Group1 and Group3. In contrast, no hybridization signals were observed in Group2 with AIMs between Group3 and Group5, suggesting that Group2 did not originate from hybridization between them (Fig. S4). A detailed examination of the distribution of each genomic group at Site6, where Group4 was collected, revealed that individuals of Group1 and Group3 were growing in close proximity (Fig. S5). These findings indicate that although Group1 and Group3 possess distinct genetic compositions, they can cross at a low frequency when they co-occur in the field.

**Figure 6.**
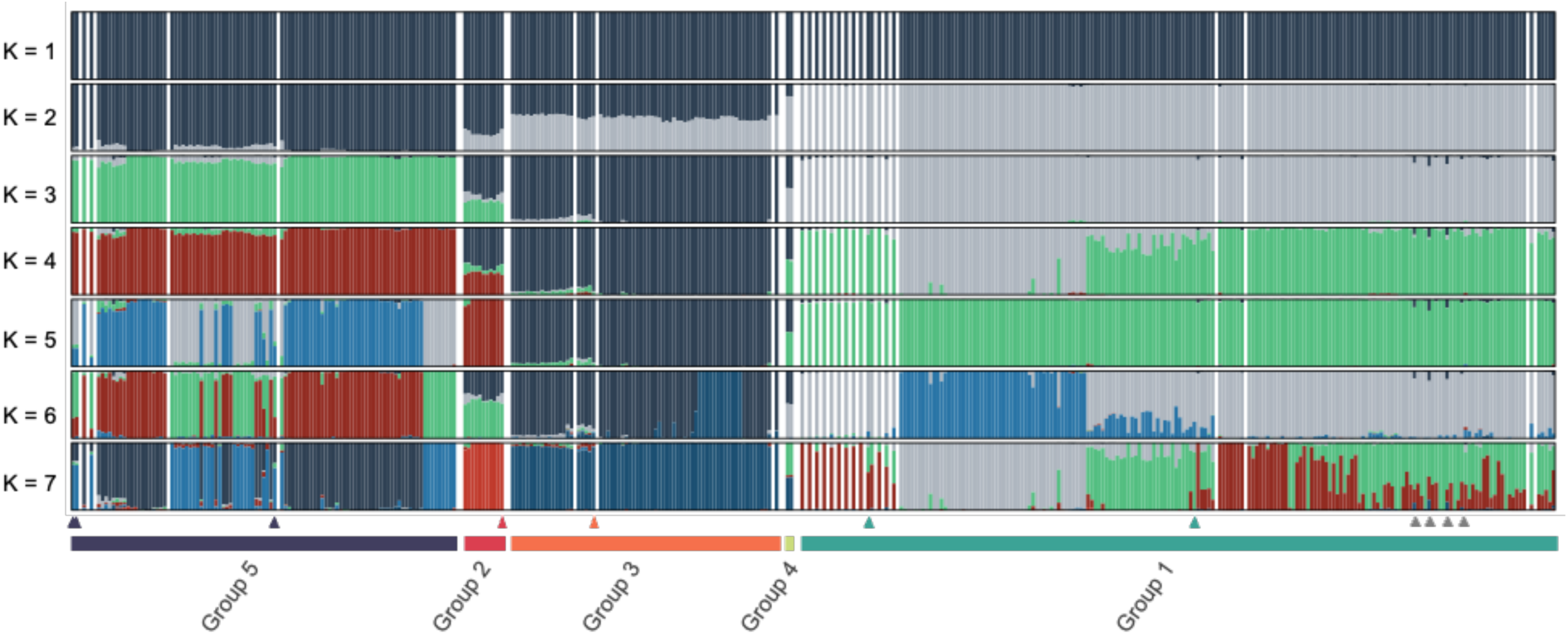
ADMIXTURE analysis of the collected samples. ADMIXTURE analysis was performed for K = 1–15 based on SNPs identified from the nuclear genome (ATBT) using the dpMIG-seq data of the collected samples. The results for K = 1–7 are shown based on the cross-validation (CV) error values (Figure. S3). In the Admixture bar plot, gaps equivalent to two samples were inserted between the genetic groups, whereas gaps equivalent to one sample were inserted according to the prefectural codes of the collected locations. Arrows at the bottom of the plot indicate the standard accessions of *E. oryzicola* (cyan), *E. crus-galli* (navy blue), and ninja (red and orange). Gray arrows indicate individuals showing partial genetic admixture between Group1 and Group3.

### 3.9 Vegetation and environmental survey at *Echinochloa* collection sites

Finally, we investigated the habitats of each *Echinochloa* genomic group. For Sites 1–6, where Groups 1–5 were found, multiple quadrats were established, and the vegetation within each quadrat and environmental variants, such as soil properties, TWI, and solar radiation were surveyed. These datasets were integrated and subjected to PCA to quantify the environmental characteristics. While the habitat of Group1 formed a cluster on the PCA plot, the habitats of Group2, Group3, and Group5 did not. These results indicate that among Japanese tetraploid *Echinochloa* species, Group2 and Group3, including ninja, inhabit a broader range of environmental niches than Group1, which includes *E. oryzicola* var. *oryzicola*.

## 4 DISCUSSIONS

### 4.1 Ninja’s positioning

Our genetic analysis revealed that ninja is a tetraploid *Echinochloa* species with an AABB-like nuclear subgenome composition, which is closely related to *E. oryzicola* var. *oryzicola* among the previously reported tetraploid species (Fig. 1, S1, 2, S2). Further population structure and phylogenetic analyses of the field-collected samples indicated that at least two types with distinct genetic compositions and evolutionary lineages exist within the ninja accessions (Fig. 4, 5, 6), and they were different from other tetraploid *Echinochloa* species worldwide (Fig. S2), suggesting that ninja are not local variants derived from previously identified tetraploid species but have their own evolutionary histories. In addition, individuals with a genetic composition similar to that of ninja were identified in samples from Hokkaido and Iwate Prefectures dating back to the 1970s (Fig. 4). These findings suggest that ninja is not an exotic species that recently invaded Japan from overseas but has been present in Japan since the 1970s.

### 4.2 Genetic and ecological diversity in tetraploid *Echinochloa* species

The genetic and ecological differences between ninja and *E. oryzicola* var. *oryzicola* suggest substantial diversity among tetraploid *Echinochloa* species in Japan. In the read-mapping analysis of pseudo-hexadecaploid genomes and PCA, ninja and *E. orzicola* var. *oryzicola* exhibited distinct read-mapping patterns and genetic compositions from one another (Fig. 1, S1, S2, 4A). Furthermore, our environmental surveys revealed that ninja inhabits a broader range of environmental niches than *E. oryzicola* var. *oryzicola* (Fig. 8). These findings suggest that the currently widespread *E. oryzicola* var. *oryzicola* individuals represent only a subset of the diversity of tetraploid *Echinochloa* species in Japan.

In our field-collected samples, Group2 (10 samples) and Group3 (67 samples), to which the ninja-standard-accessions belong, tended to be less abundant than Group1 (154 samples) and Group5 (95 samples), despite the fact that collection sites were selected from areas where ninja had previously been found (Yasuda et al., 2020). This tendency was especially pronounced in arable land. For example, Site1, Site3, and Site5, each with a high proportion of Group2 and Group3, were located in mountainous grasslands, roadside ditches, and abandoned fields, whereas Site2, Site4, and Site6, which were paddy fields or their surrounding levees, were dominated by Group1 and Group5. These observations suggest that, compared with *E. oryzicola* var. *oryzicola* and *E. crus-galli*, ninja are less adapted to contemporary agricultural practices and may have been displaced from their former habitats by human agricultural activities. Although ninja may be gradually declining in abundance and may be at risk of extinction, its presence provides a remnant of the genetic and ecological diversity of tetraploid *Echinochloa* species that existed in Japan before agricultural expansion.

### 4.3 Genetic introgression between tetraploid species

Although tetraploid ninja and *E. oryzicola* var. *oryzicola* showed a distinct genetic composition, they occurred sympatrically in paddy fields and paddy levees (Fig. S5) and hybridized to produce F_1_ individuals at a low frequency (2/333 individuals). No F_2_ individuals were detected among the collected samples (Fig. 7). However, ADMIXTURE analysis confirmed that some samples of Group1, which co-occurred with hybrid progeny, contained a small proportion of ancestral components shared with Group3 (Fig. 6). This result suggests that these samples may have arisen through backcrossing with *E. oryzicola* var. *oryzicola* following initial hybridization with ninja. Taken together, F_1_ individuals between ninja and *E. oryzicola* var. *oryzicola* have a low self-pollination efficiency, and genetic introgression between the two groups likely occurs when fertility is restored through backcrossing. A previous study also identified hybrid progeny between *E. oryzicola* var. *oryzicola* and *E. crus-galli* in field-collected samples (Wu et al., 2022), which differ in ploidy, and samples presumed to be backcrosses to the *E. crus-galli* parent have also been found (1/737 individuals). These findings confirm that *Echinochloa* species with distinct genetic backgrounds can undergo genetic mixing through crossing and subsequent backcrossing, albeit at a low frequency. Given the genetic, ecological, and morphological differences between ninja and *E. oryzicola* var. *oryzicola*, there is a possibility that novel adaptive traits related to weediness may arise in progeny as a result of their hybridization. Considering the genetic diversity of tetraploid *Echinochloa* species, including ninja, and hybridization events among them may provide new insights into the agricultural adaptation of *E. oryzicola* var. *oryzicola*.

**Figure 7.**
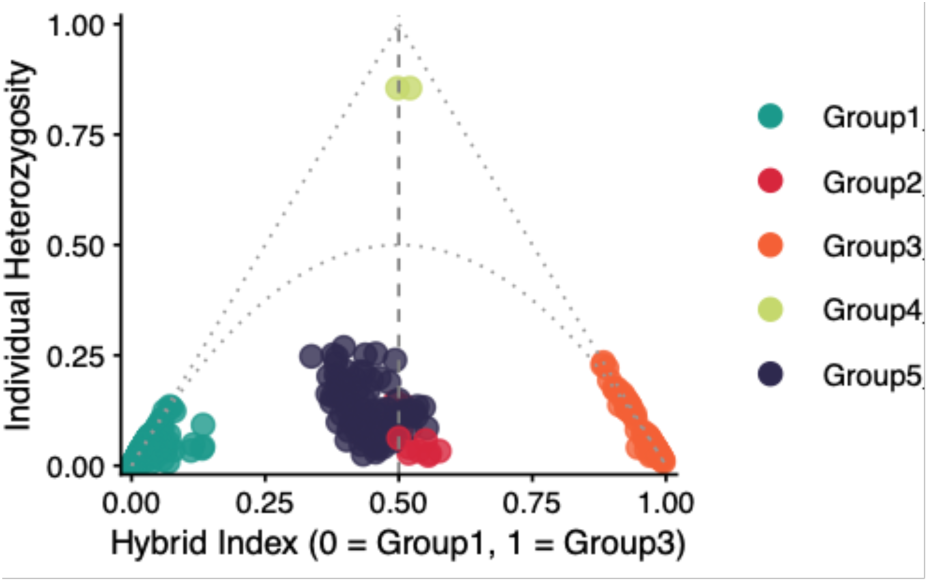
Triangle plot based on 244 ancestry-informative markers (AIMs) between Group1 and Group3. A total of 244 polymorphic sites showing fixed differences between Group1 and Group3 were selected as AIMs. The hybrid index (HI) and heterozygosity of each sample at these AIMs were analyzed and plotted as triangle plots.

### 4.4 Genetic relationship between ninja and hexaploid *Echinochloa* species

Based on the population structure analysis of the nuclear genome, genetic similarity was observed between Group2, to which ninja_1 belongs, and Group5, to which *E. crus-galli* belongs (Fig 6), and it was inferred that this genetic similarity did not arise from recent hybridization (Fig. S4). Our observation of the overlap of their habitats (Fig. 8) also suggests the possibility that not the paddy field-adapted *E. oryzicola* var. oryzicola, but a dry-environment-adapted tetraploid species, such as ninja, was a direct ancestor of hexaploid *E. crus-galli*. In addition, phylogenetic analysis of chloroplast genomes revealed a close relationship between the Chinese and Malaysian accessions of *E. crus-galli* var*. crus-galli* and ninja_1 in ECpG_clade1 (Fig. 2A). In Japan, several accessions of *E. crus-galli* var. *formosensis* have also been reported to possess chloroplast genomes closely related to those of *E. oryzicola* accessions (Yasuda et al., 2020), which, based on the phylogenetic relationships observed in this study, are likely to correspond to ECpG_clade1. Although we cannot exclude the possibility that *E. crus-galli* in ECpG_clade1 originated through chloroplast capture, given the limited number of *E. crus-galli* accessions in ECpG_clade1 included in the global analysis to date (Wu et al., 2022), if distinct allopolyploidization events are involved in *E. crus-galli* origination, it will be necessary to investigate the possibility that dry-environment-adapted tetraploid *Echinochloa*, such as ninja, was involved as one of the parental species that produced *E. crus-galli* with an ECpG_clade1-type chloroplast.

**Figure 8.**
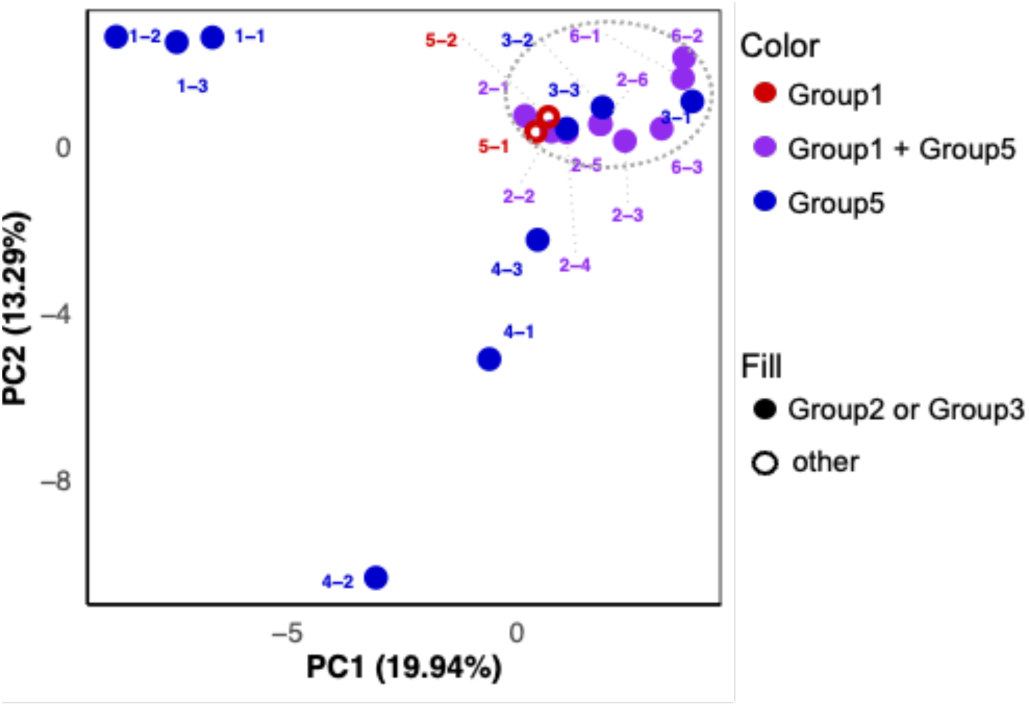
PCA of vegetation and environmental conditions across the habitats of *Echinohloa* Groups 1–5. The PCA integrated vegetation composition, soil properties, TWI, and potential incoming solar radiation at Sites 1–6 (Fig. 4B) to characterize habitat conditions associated with each genetic group. Each plot is colored according to the presence or absence of Group1 and Group5 at each site, whereas sites where Group2 and Group3 were collected are indicated by filled plots. The dotted grey circle indicates the habitat conditions associated with Group1, which includes E. oryzicola var. oryzicola. The quadrat numbers within each survey site are shown next to the plots.

In addition, the ECpG_clade2-type chloroplast of *E. crus-galli*, traditionally considered to be derived from the C subgenome donor (Aoki & Yamaguchi, 2008), may instead have originated from tetraploid species with an AABB-like subgenome composition in ECpG_clade2, such as *E. walteri* or *E. oryzicola* var. *hainanensis* (Figs. 1, 2). This interpretation is more consistent with the patterns observed in the phylogenies of the nuclear and chloroplast genomes of global *Echinochloa* samples. Based on the divergence-time analysis of *Echinochloa* genomes (Wu et al., 2022; Ye et al., 2020), the nuclear genome of diploid *E. haploclada* appears to have diverged most closely from subgenome C of *E. crus-galli*. If the chloroplast genome of *E. crus-galli* originated from a diploid parent carrying subgenome C, the chloroplast genomes of *E. crus-galli* should be most closely related to those of *E. haploclada,* which possesses a genome composition closely related to subgenome C. However, recent chloroplast phylogenies, including diverse tetraploid species (Wu et al., 2022; Yasuda & Nakayama, 2019), as well as our analysis including ninja, consistently showed that the chloroplast genome of *E. crus-galli* was more closely related to those of tetraploid species with AABB-like subgenome compositions, such as *E. walteri* and *E. oryzicola* var. *hainanensis* than *E. haploclada* in ECpG_clade2 (Fig. 1A). These findings suggest that *E. crus-galli* in ECpG_clade2 may have also originated from a tetraploid species other than *E. oryzicola* var. *oryzicola*.

### 4.5 Morphological and ecological traits differences

In this study, we investigated the genetic relationships and distribution of ninja. However, we have not evaluated their detailed morphological or ecological characteristics. As reported in previous studies (Yasuda et al., 2020), unlike the closely related tetraploid *E. oryzicola* var. *oryzicola*, ninja has semi-erect tillers similar to those of hexaploid *E. crus-galli*. In addition, for spikelet size, diversity has been reported among ninja ranging from as small as *E. crus-galli* to as large as *E. oryzicola* var. *oryzicola*. In this study, individuals with deep purple leaf coloration, as seen in *E. crus-galli*, were also observed (Figure S.6). Ninja exhibit diversity in various morphological traits, but these have not been quantified. Furthermore, differences in the habitats of ninja_1 and ninja_2 have been observed, with ninja_1 often being distributed sympatrically with *E. crus-galli* in grasslands or vacant lots, whereas ninja_2 was distributed sympatrically with *E. oryzicola* var. *oryzicola* on paddy levees (Fig. 4, S5). Based on this observation, ninja individuals are also thought to exhibit diverse degrees of flood adaptation, but the details remain unknown. If the adaptive traits of ninja and their similarities or differences with other *Echinochloa* species are clarified, it will be possible to interpret the adaptive evolution of *Echinochloa* species with their diverse genetic backgrounds. Future research should investigate the morphological and ecological characteristics of ninja through common garden cultivation or reciprocal transplant experiments.

### 4.6 The unclassified cryptic *Echinochloa* species in Japan

The authors used the term “ninja” not for a specific *Echinochloa* species but as a general term for unclassified cryptic tetraploid *Echinochloa* plants found in Japan (Yasuda et al., 2020). To avoid confusion in nomenclature, we did not determine the taxonomic position of ninja in this study. Although, in recent years, foundational information on the genetic diversity of *Echinochloa* at national and global scales has been established (Sato et al., 2023; Wu et al., 2022; Ye et al., 2019), no study has yet been conducted to evaluate the genetic diversity of *Echinochloa* plants across Japan, and it seems premature to determine the taxonomic position of ninja based on our limited samples from northern Japan. If our genetic approach is implemented and the survey area is further expanded, it is highly likely that previously unrecognized cryptic species, such as the ninja, will be discovered. In the future, if the genetic, ecological, and morphological diversity of *Echinochloa* species throughout Japan can be evaluated with the diversity of global samples, it should be possible to establish the taxonomic position of each unclassified sample, including ninja. The comprehensive discovery and analysis of diverse *Echinochloa species* in Japan is an important research topic.

## ACKNOWLEDGMENTS

We thank Norihiro Shimizu for collecting the *Echinochloa* plants in Japan in the 1970s. We also thank the NARO Genebank (Genetic Resources Research Center, National Agriculture and Food Research Organization, Japan) for maintaining and providing these resources. This work was supported by the Japan Society for the Promotion of Science (JSPS) Grant-in-Aid for JSPS Fellows (Grant Number: 23KJ1360).

## AUTHOR CONTRIBUTIONS

Tomomi Kubo conceived and planned the study, conducted field surveys, collected and analyzed the data, interpreted the results, and wrote the manuscript. Hikari Ikeda participated in field vegetation and environmental surveys and conducted the corresponding analyses. Kentaro Yasuda and Shunji Kurokawa participated in field sampling and vegetation surveys. They also contributed to the study planning, interpretation of the results, and manuscript preparation through extensive discussions with the first author.

## DISCLOSURE STATEMENT

The authors declare no conflict of interest.

## DATA AVAILABILITY

The dpMIG-seq data were deposited with links to the BioProject accession number PRJDB43033 in the DDBJ BioProject database. Samples with unsuccessful DNA extraction and those with substantially lower read counts were excluded from the deposited dataset.

## Notes

### Competing Interest Statement

The authors have declared no competing interest.

