## Supplementary Materials for "Genomic and ecological characterization of unclassified tetraploid *Echinochloa* plants distributed in northern Japan"

**SUPPORTING INFORMATION**

Additional Supporting Information is available in the online version of this article at the publisher's website.

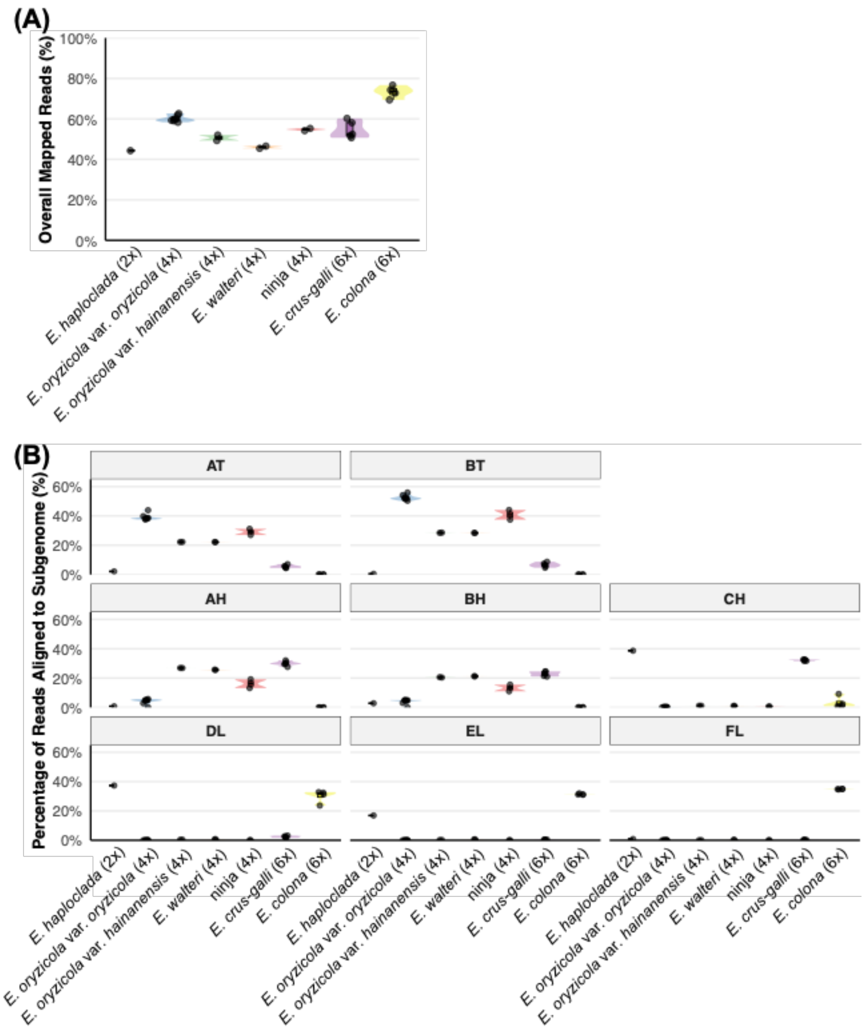


**Figure S1. Read mapping ratio to the pseudo-hexadecaploid genome**
**A: Overall read mapping rates (%)**
**B: Subgenome-specific read mapping proportions (%)**

The resequencing reads of *Echinochloa species,* including ninja, were mapped to a pseudo-hexadecaploid genome comprising ATBT, AHBHCH, and DHEHFH. Overall read mapping rates and subgenome-specific read-mapping proportions were compared among the species.

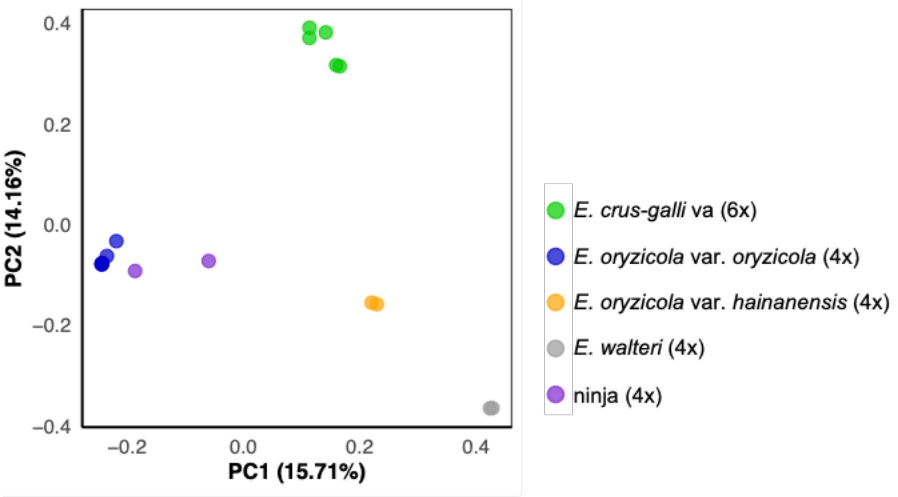


**Figure S2. PCA based on the nuclear genome-derived SNPs**

The resequencing reads of *Echinochloa species,* including ninja, were mapped to the nuclear genome of *E. oryzicola*, and the resulting SNPs were used for PCA.

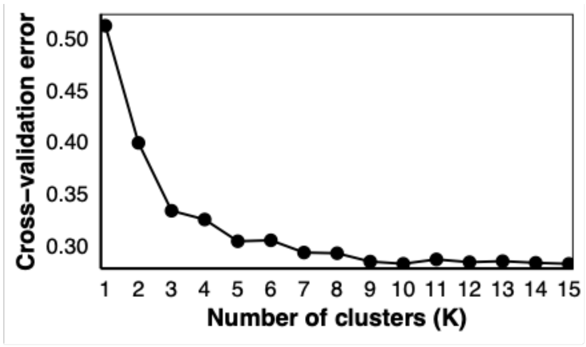


**Figure S3. Cross-validation (CV) error values from ADMIXTURE analysis for K = 1–15.**

ADMIXTURE analysis was performed based on nuclear (ATBT)-derived SNPs using dpMIG-seq datasets of the collected samples. The CV error values estimated for each K value are shown.

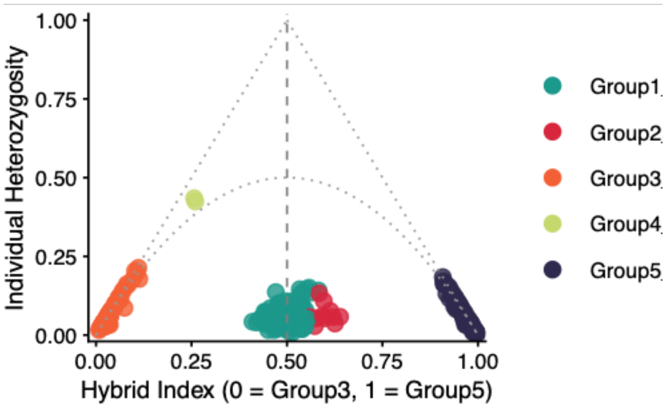

**Figure S4. Triangle plot based on 272 ancestry-informative markers (AIMs) between Group3 and Group5.**

A total of 272 polymorphic sites showing fixed differences between Group3 and Group5 were selected as AIMs. The hybrid index (HI) and heterozygosity of each sample at these AIMs were analyzed and plotted as triangle plots.

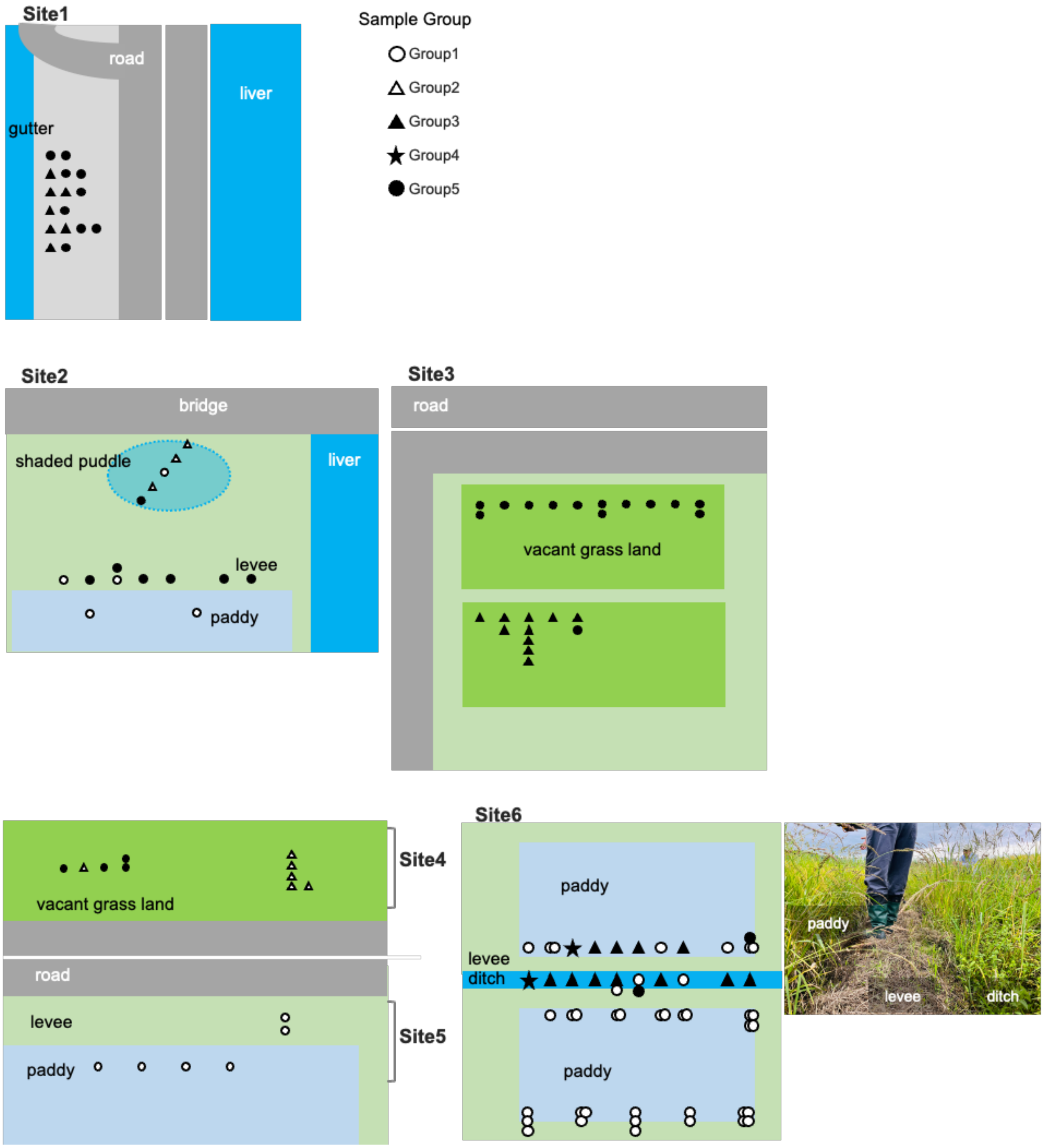


**Figure S5. A Picture and schematic representations of Sites 1–6.**

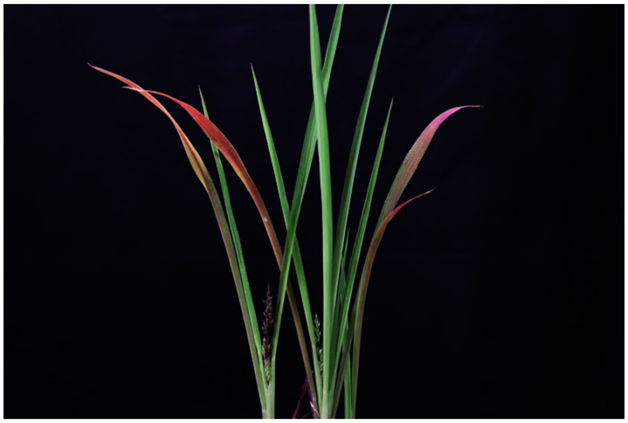

**Figure S6. Picture of an individual ninja with a purple leaf.**
